# Modern human changes in regulatory regions implicated in cortical development

**DOI:** 10.1101/713891

**Authors:** Juan Moriano, Cedric Boeckx

## Abstract

Recent paleogenomic studies have highlighted a very small set of proteins carrying modern human-specific missense changes in comparison to our closest extinct relatives. Despite being frequently alluded to as highly relevant, species-specific differences in regulatory regions remain understudied. Here, we integrate data from paleogenomics, chromatin modification and physical interaction, and single-cell gene expression of neural progenitor cells to report a set of genes whose enhancers and/or promoters harbor modern human single nucleotide changes that appeared after the split from the Neanderthal/Denisovan lineage. These regulatory regions exert their functions at early stages of cortical development and control a set of genes among which those related to chromatin regulation stand out. This functional category has not yet figured prominently in modern human evolution studies. Specifically, we find an enrichment for the SETD1A histone methyltransferase complex, known to regulate WNT-signaling for the generation and proliferation of intermediate progenitor cells.

## 1 Introduction

Progress in the field of paleogenomics has allowed researchers to study the genetic basis of modern human-specific traits in comparison to our closest extinct relatives, the Neanderthals and Denisovans [1]. One such trait concerns the period of growth and maturation of the brain, which is a major factor underlying the characteristic ‘globular’ head shape of modern humans [2]. Comparative genomic analyses using high-quality Neanderthal/Denisovan genomes [3–5] have revealed missense changes in the modern human lineage affecting proteins involved in the division of neural progenitor cells, key for the proper generation of neurons in an orderly spatiotemporal manner [4, 6]. But the total number of fixed missense changes amounts to less than one hundred proteins [1, 6]. This suggests that changes falling outside protein-coding regions may be equally relevant to understand the genetic basis of modern human-specific traits, as proposed more than four decades ago [7]. In this context it is noteworthy that human positively-selected genomic regions were found to be enriched in regulatory regions [8], and that signals of negative selection against Neanderthal DNA introgression were reported in promoters and conserved genomic regions [9].

Here, we report a set of genes under the control of regulatory regions that harbor modern human-lineage genetic changes and are active at early stages of cortical development (Figure 1). We integrated data on chromatin immunoprecipitation and open chromatin regions identifying enhancers and promoters active during human cortical development, and the genes regulated by them as revealed by chromatin physical interaction data, together with paleogenomic data of single-nucleotide changes (SNC) distinguishing modern humans and Neanderthal/Denisovan lineages. This allowed us to uncover those enhancer and promoters that harbor modern human SNC (thereafter, mSNC) at fixed or nearly fixed frequency (as defined by [6]) in present-day human populations and where the Neanderthals/Denisovans carry the the ancestral allele (*Methods* section 4.1). Next, we analysed single-cell gene expression data and performed co-expression network analysis to identify the genes plausibly under human-specific regulation within genetic networks in neural progenitor cells (*Methods* sections 4.2-4.3). Many of the genes controlled by regulatory regions satisfying the aforementioned criteria are involved in chromatin regulation, and prominently among these, the SETD1A histone methyltransferase (SETD1A/HMT) complex. This complex, which has not figured prominently in the modern human evolution literature until now, appears to have been targeted in modern human evolution and specifically regulates the indirect mode of neurogenesis through the control of WNT/*β*-CATENIN signaling.

**Figure 1:**
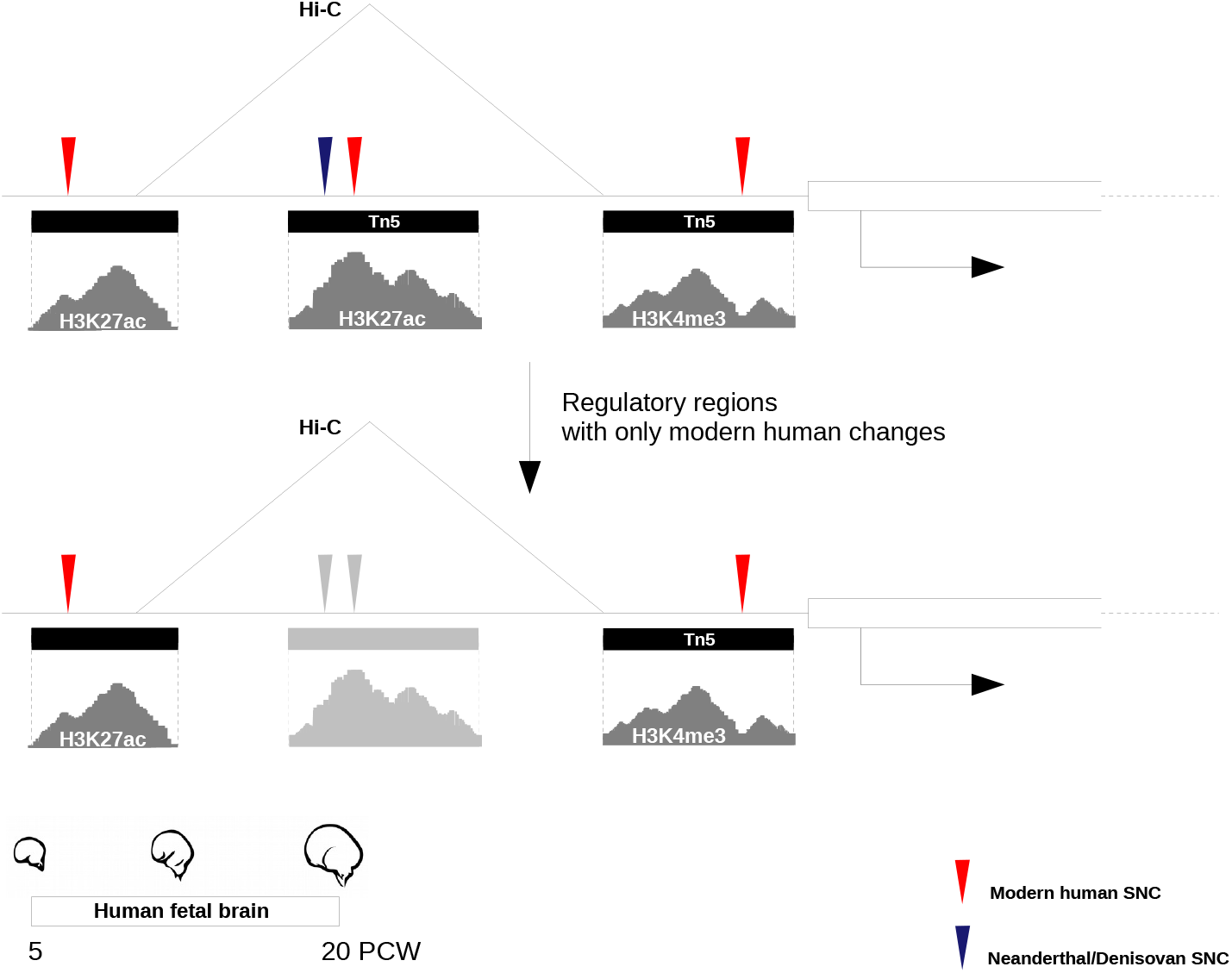
Regulatory regions characterized in this study. Active enhancers are typically located in regions of open chromatin and nucleosomes in their vicinity are marked by histone modifications H3K27 acetylation and H3K4 mono-methylation. By contrast, H3K4 tri-methylation defines active promoters [57]. We considered signals of active enhancers and promoters, as well as transposase (Tn5)-accessible chromatin regions, in the developing human brain (from 5 to 20 post-conception weeks) that harbor modern human single-nucleotide changes filtering out those regulatory regions that also contain Neanderthal/Denisovan changes. Chromosome conformation capture (Hi-C) data revealed the genes controlled by these regulatory regions.

## 2 Results

212 genes were found associated to regulatory regions active in the developing human cortex (from 5 to 20 post-conception weeks) that harbor mSNCs and do not contain Neanderthal/Denisovan changes (Suppl. Mat. Tables S1 & S2). Among these, some well-studied disease-relevant genes are found: *HTT* (Huntington disease) [10], *FOXP2* (language impairment) [11], *CHD8* and *CPEB4* (autism spectrum disorder) [12, 13], *TCF4* (Pitt-Hopkins syndrome and schizophrenia) [14, 15], *GLI3* (macrocephaly and Greig cephalopolysyndactyly syndrome) [16], *PHC1* (primary, autosomal recessive, microcephaly-11) [17], *RCAN1* (Down syndrome) [18], and *DYNC1H1* (cortical malformations and microcephaly) [19].

Twelve out of the 212 genes contain fixed mSNCs in enhancers (*NEUROD6, GRIN2B, LRRC23, RNF44, KCNA3, TCF25, TMLHE, GLI4, DDX12P, PLP2, TFE3, SPG7*), with *LRRC23* having three such changes, and *GRIN2B, DDX12P* and *TFE3*, two each. Fourteen genes have fixed mSNCs in their promoters (*LRRC23, SETD1A, FOXJ2, LIMCH1, ZFAT, SPOP, DLGAP4, HS6ST2, UBE2A, FKBP1A, RPL6, LINC01159, RBM4B, NFIB*). Only one gene, *LRRC23*, exhibits fixed changes in both its enhancer and promoter regions. To identify putatively mSNC-enriched regions, we ranked regulatory regions by mutation density (*Methods* section 4.4). Top candidates enhancers (top 5% in hits-per-region length distribution) were associated with potassium channel KCNQ5, actin-binding protein FSCN1, and neuronal marker NEUROD6. Top candidate promoters were linked to cytoplasmic dynein DYNC1H1, nuclear factor NFIB, PHD and RING finger domains-containin PHRF1, and kinesin light KLC1 (Suppl. Mat. Table S3 & S4). Interestingly, most of these are known to be involved in later stages of neurogenesis (differentiation and migration steps).

A significant over-representation was found for enhancers (permutation test; p-value 0.01) and promoters (permutation test; p-value 10^*−*4^) overlapping with putative modern human positively-selected regions [8]. In addition, we found a significant enrichment for enhancers (permutation test; p-value 0.04; while for promoter regions p-value 0.08) overlapping with genetic loci associated to schizophrenia [20]. By contrast, no significant overlap was found for enhancers/promoters and autism spectrum disorder risk variants ([21], retrieved from [22]) (Suppl. Fig. S1). We also performed motif enrichment analysis for our enhancer/promoter region datasets (*Methods* section 4.4). We found a motif enrichment in enhancer regions for transcriptional regulators IRF8, PU.1, CTCF (Benjamini q-value 0.01) and OCT4 (Benjamini q-value 0.02); while for promoter regions a motif enrichment was detected for the zinc finger-containing (and WNT signaling regulator) ZBTB33 (Benjamini q-value 0.03).

Next, we evaluated relevant gene ontology and biological categories in our 212 gene list (*Methods* section 4.4). We identified a substantial proportion of genes related to beta-catenin binding (GO:0008013; h.t.: adj p-value 0.11) and transcriptional regulation (GO:0044212; hypergeometric test (h.t.): adj p-value 0.17), and detected a significant enrichment from the CORUM protein complexes database for the SETD1A/HMT complex (CORUM:2731; h.t.: adj p-value 0.01). Indeed, three members of the SETD1A/HMT complex are present in our 212 gene list: SETD1A (fixed mSNC in promoter), ASH2L (mSNC in enhancer) and WDR82 (mSNC in enhancer). SETD1A associates to the core of an H3K4 methyltransferase complex (ASH2L, WDR5, RBBP5, DPY30) and to WDR82, which recruits RNA polymerase II, to promote transcription of target genes through histone modification H3K4me3 [23]. Furthermore, the *SETD1A* promoter and the *WDR82* enhancer containing the relevant changes fall within putative positively-selected regions in the modern human lineage [8].

The abundance of transcriptional regulators and the specific enrichment for the SETD1A/HMT led us to examine the gene expression programs likely under their influence in neural progenitor cells. From 5 to 20 post-conception weeks, different types of cells populate the germinal zones of the developing cortex (Figure 2). We re-analyzed gene expression data at single-cell resolution from a total of 762 cells from the developing human cortex, controlling for cell-cycle heterogeneity as a confounding factor in the analysis of progenitor populations (*Methods* section 4.2). We focused on two progenitor cell-types—radial glial and intermediate progenitor cells (RGCs and IPCs, respectively)—two of the main types of progenitor cells that give rise, in an orderly manner, to the neurons present in the adult brain (Figure 2). Two sub-populations of RGCs were identified (*PAX6* +/*EOMES*-cells), and three sub-populations of IPCs were detected (*EOMES*-expressing cells, with cells retaining *PAX6* expression and some expressing differentiation marker *TUJ1*), largely replicating what has been reported in the original publication for this dataset (Suppl. Fig. 2). We next identified genetic networks (based on highly-correlated gene expression levels) in the different sub-populations of progenitor cells (except for IPC sub-population 3, which was excluded due to the low number of cells) (*Methods* section 4.3; Suppl. Fig. S3 & S4).

**Figure 2:**
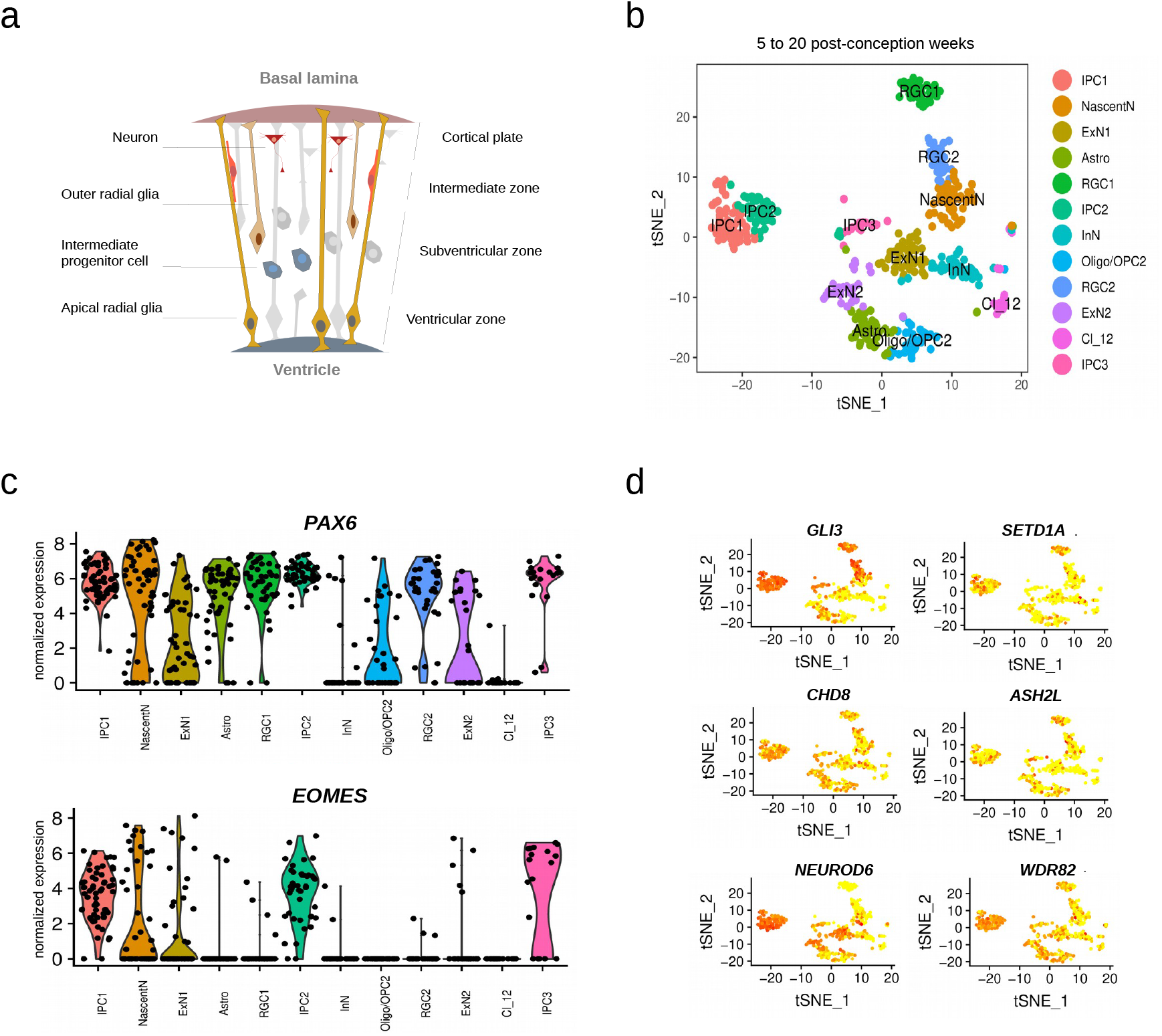
Cell-type populations at early stages of cortical development. (a) Apical radial glial cells (RGCs) populate the ventricular zone and prolong one process apically to the ventricular surface and another one to the basal lamina, which serves as a scaffold for neuronal migration. RGCs also proliferate and differentiate to give rise to another RGC, a basal intermediate progenitor (indirect neurogenesis), or a neuron (direct neurogenesis) [58]. Intermediate progenitor cells (IPCs) are basal progenitors lacking of apical-basal cell polarity. IPCs migrate to the subventricular zone and, after a couple of self-renewal divisions, differentiate to give rise to two neurons [58]. (b) The tSNE plot shows twelve clusters detected analyzing a total of 762 cells. (c) The violin plots show expression of two markers (*PAX6, EOMES*) across the different clusters, distinguishing between RGCs and IPCs. (d) The miniature tSNE plots show the distribution across the sub-populations of a selection of genes discussed in the main text. IPC: Intermediate progenitor cells; NascentN: Nascent neurons; ExN: Excitatory neurons; Astro: Astrocytes; RGC: Radial glial cells; InN: Interneurons; Oligodendrocyte progenitor cells: OPC; Oligo: Oligodendrocytes; Cl 12: Cluster 12 (unidentified cells).

An over-representation of genes related to the human phenotype ontology term ‘Neurodevelopmental abnormality’ was detected in the RGC-2 turquoise module (HP:0012759; h.t.: adj p-value 0.03, Suppl. Mat. Table S5). Indeed, a considerable amount of genes were found to be associated to phenotype terms ‘Neurodevelopmental delay’ and ‘Skull size’ (HP:0012758 and HP:0000240, respectively; h.t.: adj p-value 0.07 and 0.13, respectively; Suppl. Mat. Table S6). Two chromatin regulators with mSNC in regulatory regions are present in these two ontology terms and are associated to neurodevelopmental disorders: KDM6A (mSNC in promoter), which associates to the H3K4 methyltransferase complex [23], and is mutated in patients with Kabuki syndrome [24]; and PHC1 (mSNC in promoter), a component of the repressive complex PRC1 [23], found in patients with primary microcephaly-11 [17]. Among the total genes related to the ‘Skull size’ term (n = 109), we found an over-representation of genes (*CDON, GLI3, KIF7, GAS1*) related to the hedgehog signaling pathway (KEGG:04340; h.t.: adj p-value 0.05). Of these, *GLI3* (mSNC in promoter) is perhaps the most salient member. *GLI3* is a gene linked to macrocephaly [16] and under putative modern human positive selection [8]. Considering that hedgehog signaling plays a critical role in basal progenitor expansion [25], we note the presence in this turquoise module of the outer radial glia-specific genes *IL6ST* and *STAT3* [26]. The forkhead-box transcription factor FOXP2 is also present in RGC-2 turquoise module and associated to the ‘Neurodevelopmental delay’ ontology term. Its promoter harbors an almost fixed (*>*99%) mSNC. FOXP2 is a highly conserved protein involved in language-related disorders whose evolutionary changes are particularly relevant for understanding human cognitive traits [27]. This mSNC (7:113727420) in the *FOXP2* promoter adds new evidence for a putative modern human-specific regulation of *FOXP2* together with the nearly fixed intronic SNC that affects a transcription factor-binding site [27].

While we did not detect a specific enrichment in the modules containing SETD1A/HMT complex components ASH2L or WDR82 genes, the IPC-2 midnightblue module, which contains SETD1A, shows an enrichment for a *β*-CATENIN-containing complex (SETD7-YAP-AXIN1-*β*-CATENIN complex; CORUM:6343; h.t.: adj p-value 0.05; Suppl. Mat. Table S7) and indeed contains WNT-effector TCF3, which harbors nearly fixed missense mutations in modern humans [6]. SETD1A is known to interact with *β*-CATENIN [28, 29] and increase its expression to promote neural progenitor proliferation [30].

## 3 Discussion

By integrating data from paleogenomics and chromatin interaction and modification, we identified a set of genes controlled by regulatory regions that are active during early cortical development and contain single nucleotide changes that appeared in the modern human lineage after the split from the Neanderthal/Denisovan lineage. This study complements previous research focused on protein-coding changes [4, 6] and helps extend the investigation of species-specific differences in cortical development that has so far relied on detailed comparisons between humans and non-human primates [31–35].

The regulatory regions reported here significantly overlap with putative modern human positively-selected regions and schizophrenia genomic loci, and control a set of genes among which we find a high number related to chromatin regulation, and most specifically the SETD1A/HMT complex. Regulators of chromatin dynamics are known to play key roles during cell-fate decisions through the control of specific transcriptional programs [36–38]. Both SETD1A and ASH2L, core components of the HMT complex, regulate WNT/*β*-CATENIN signaling [28–30, 39], which influences cell-fate decisions by promoting either self-maintenance or differentiation depending on the stage of progenitor differentiation (Figure 3).

**Figure 3:**
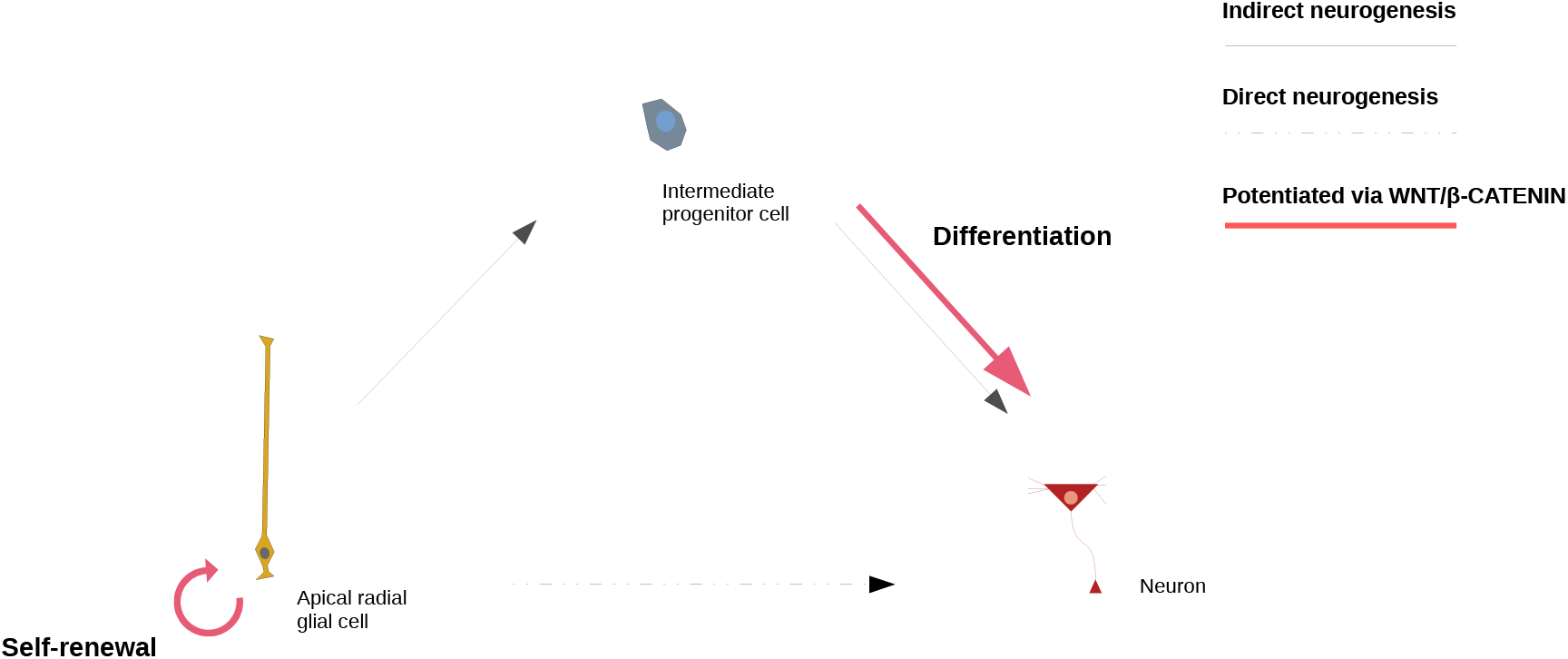
Progenitor cell-fate decisions shaped by WNT/*β*-CATENIN signaling. Based on studies in mice, it is hypothesized that early during neurodevelopment, WNT/*β*-CATENIN signaling promotes neural stem and progenitor cell self-renewal whereas its depletion causes premature neuronal differentiation [59–61]; later on, its down-regulation is required for generation of intermediate progenitor cells from radial glial cells [61, 62]. Lastly, WNT/*β*-CATENIN signaling promotes differentiation of intermediate progenitors to produce neurons [60].

SETD1A (fixed mSNC in promoter), implicated in schizophrenia and developmental language impairment [40, 41], acts in collaboration with a histone chaperone to promote proliferation of neural progenitor cells through H3K4 trimethylation at the promoter of *β*-CATENIN, while its knockdown causes reduction in proliferative neural progenitor cells and an increase in cells at the cortical plate [30]. In addition, one of SETD1A direct targets is the WNT-effector TCF4 [42], whose promoter also harbors a mSNC. Similarly, ASH2L specifically regulates WNT signaling: Conditional knock-out of *ASH2L* significantly compromises the proliferative capacity of RGCs and IPCs by the time of generation of upper-layer neurons, with these progenitor cells showing a marked reduction in H3K4me3 levels and downregulation of WNT/*β*-CATENIN signaling-related genes (defects that can be rescued by over-expression of *β*-CATENIN) [39]. Taken together, depletion of components of the SETD1A/HMT complex impairs the proliferative capacity of progenitor cells, altering the indirect mode of neurogenesis, with a specific regulation of the conserved WNT signaling. Our data points toward a putative modern human positive selection of their regulatory regions. Interestingly, in a recent work studying species-specific differences in chromatin accessibility using brain organoids, regulatory regions associated to *SETD1A* and *WDR82* were found in differentially-accessible chromatin regions in human organoids in comparison to chimpanzee organoids, with the *SETD1A* region overlapping with a human-gained histone modification signal when compared to macaques [35].

The dysfunction of chromatin regulators is among the most salient features behind causative mutations in neurodevelopmental disorders [43]. Our data highlights chromatin modifiers and remodelers that play prominent roles in neurodevelopmental disorders affecting brain growth and facial features. Along with the aforementioned chromatin regulators PHC1 (microcephaly) and KDM6A (Kabuki syndrome), another paradigmatic example is the ATP-dependent chromatin remodeler CHD8 (mSNC in enhancer), which controls neural progenitor cell proliferation through WNT-signaling related genes [44, 45]. CHD8 is a high-risk factor for autism spectrum disorder and patients with CHD8 mutations characteristically present macrocephaly and distinctive facial features [12]. Intriguingly, another ATP-dependent chromatin remodeler, CHD2 (mSNC in enhancer), presents a motif in the SETD1A promoter region containing the fixed mSNC (16:30969654; UCSC Genome Browser). The study of modern human evolutionary changes affecting chromatin regulators integrated with the examination of neurodevelopmental disorders promises to improve our understanding of modern human-specific brain ontogenetic trajectories.

We have focused on the early stages of cortical development. While single-cell gene expression data of neural progenitor cells still remains limited, future integration of these data with other datasets covering different neocortical regions [46] will shed further light on modern human changes and cortical areas-specific progenitor cells. We acknowledge, in addition, that the genetic changes distinguishing modern humans and Neanderthals/Denisovans may be relevant at other stages of neurodevelopment, including the adult human brain. Progress in single-cell multi-omic technologies applied to brain organoid research will be critical to assess the impact of such changes in the diverse neural and non-neural cell-types through different developmental stages. Moreover, we excluded the examination of regulatory regions harboring Neanderthal/Denisovan changes due to the low number of high-quality genomes from Neanderthal/Denisovan individuals, which makes the determination of allele frequency in these species unreliable. We hope that the availability of a higher number of high-quality genomes for these species in the future will make such examination feasible.

## 4 Methods

### 4.1 Data processing

Integration and processing of data from different sources was performed using IPython v5.7.0. We used publicly available data from [6] of SNC in the modern human lineage (at fixed or above 90% frequency in present-day human populations) and Neanderthal/Denisovan changes. [6] analyzed high-coverage genotypes from one Denisovan and two Neanderthal individuals to report a catalog of SNC that appeared in the modern human lineage after their split from Neanderthals/Denisovans. Similarly, [6] also reported a list of SNC present in the Neanderthal/Denisovan lineages where modern humans carry the inferred ancestral allele.

For enhancer–promoter linkages, we used publicly available data from [47], based on transposase-accessible chromatin coupled to sequencing and integrated with chromatin capture via Hi-C data, from 15 to 17 post-conception weeks of the developing human cortex. A total of 92 promoters and 113 enhancers were selected as harboring mSNC and beind depleted of Neanderthal/Denisovan SNC (from a total of 2574 enhancers and 1553 promoters present in the original dataset). Additionally, we completed the previous dataset filtering annotated enhancer-gene linkages via Hi-C from the adult prefrontal cortex [48] (PsychENCODE resource portal: http://resource.psychencode.org/). In this case, enhancers (n = 32803) were selected for further analyses if their coordinates completely overlapped with signals of active enhancers (H3K27ac) (that do not overlap with promoter signals (H3K4me3)) from the developing human cortex between 7 to 12 PCW [49]. A total of 43 enhancers, containing mSNC but free of Neanderthal/Denisovan SNC, passed this filtering. As a whole, the final integrated dataset covered regulatory regions active at early stages of human prenatal cortical development and linked to 212 genes. The coordinates (hg19 version) of the regulatory regions containing mSNC are available in the Supplementary Material Tables S1 & S2.

Human positively-selected regions coordinates were retrieved from [8].

### 4.2 Single-cell RNA-seq analysis

The single-cell transcriptomic analysis was performed using the Seurat package v2.4 [50] in RStudio v1.1.463 (server mode).

Single-cell gene expression data was retrieved from [49] from PsychENCODE portal (http://development.psychencode.org/#). We used raw gene counts thresholding for cells with a minimum of 500 genes detected and for genes present at least in 10% of the total cells (n=762). Data was normalized using ”LogNormalize” method with a scale factor of 1000000. We regressed out cell-to-cell variation due to mitochondrial and cell-cycle genes (*ScaleData* function). For the latter, we used a list of genes ([51]) that assigns scores genes to either G1/S or G2/M phase (function *CellCycleScoring*), allowing us to reduce heterogeneity due to differences in cell-cycle phases. We further filtered cells (*FilterCells* function) setting a low threshold of 2000 and a high threshold of 9000 gene counts per cell, and a high threshold of 5% of the total gene counts for mitochondrial genes.

We assigned the label ‘highly variable’ to genes whose average expression value was between 0.5 and 8, and variance-to-mean expression level ratio between 0.5 and 5 (*FindVariableGenes* function).

We obtained a total of 4261 genes for this category. Next, we performed a principal component analysis on highly variable genes and determined significance by a jackStraw analysis (*JackStraw* function). We used the first most significant principal components (n = 13) for clustering analysis (*FindClusters* function; resolution = 3). Data was represented in two dimensions using t-distributed stochastic neighbor embedding (*RunTSNE* function). The resulting twelve clusters were plotted using *tSNEplot* function. Cell-type assignment was based on the metadata from the original publication [49].

### 4.3 Weighted gene co-expression network analysis

For the gene co-expression network analysis we used the WGCNA R package [52, 53]. For each population of progenitor cells (RGC-1, 34 cells (15017 genes); RGC-2, 30 cells (14747 genes); IPC-1, 52 cells (15790 genes); IPC-2, 41 cells (15721 genes); IPC3 population was excluded due to low number of cells), log-transformed values of gene expression data were used as input for weighted gene co-expression network analysis. A soft threshold power was chosen (12, 12, 14, 12 for RGC-1, RGC-2, IPC-1, IPC-2 populations, with R2: 0.962, 0.817, 0.961, 0.918, respectively) and a bi-weight mid-correlation applied to compute a signed weighted adjacency matrix, transformed later into a topological overlap matrix. Module detection (minimum size 200 genes) was performed using function *cutreeDynamic* (method = ’hybrid’, deepSplit = 2), getting a total of 32, 26, 9, 23 modules for RGC-1, RGC-2, IPC-1, IPC-2, respectively (Suppl. Fig. S3 & S4).

### 4.4 Enrichment analysis

We ranked regulatory regions by mutation density calculating number of single nucleotide changes per regulatory region length (for those regions spanning at least 1000 base pairs). Top candidates were those raking in the distribution within the 5% out of the total number of enhancers or promoters (Suppl. Mat Tables S3 & S4). The g:Profiler2 R package [54] was used to perform enrichment analyses (hypergeometric test; correction method ‘gSCS’) for gene/phenotype ontology categories, biological pathways (KEGG, Reactome) and protein databases (CORUM, Human Protein Atlas) for the gene lists generated in this study. Permutation tests (10,000 permutations) were performed to evaluate enrichment of enhancers/promoters regions in different genomic regions datasets using the R package regioneR [55]. The Hypergeometric Optimization of Motif EnRichment (HOMER) software v4.10 [56] was employed for motif discovery analysis, selecting best matches (Benjamini q-value *<* 0.05) of known motifs (n = 428; ChIP-seq-based) in our promoter and enhancer datasets.

## Author contributions

Conceptualization: C.B. & J.M.; Data Curation: J.M.; Formal Analysis: J.M.; Funding Acquisition: C.B.; Investigation: C.B. & J.M.; Methodology: C.B. & J.M.; Software: J.M.; Supervision: C.B.; Visualization: C.B. & J.M.; Writing – Original Draft Preparation: C.B. & J.M.; Writing – Review & Editing: C.B. & J.M.

## Funding

C.B. acknowledges research funds from the Spanish Ministry of Economy and Competitiveness/FEDER (grant FFI2016-78034-C2-1-P), Marie Curie International Reintegration Grant from the European Union (PIRG-GA-2009-256413), research funds from the Fundació Bosch i Gimpera, MEXT/JSPS Grant-in-Aid for Scientific Research on Innovative Areas 4903 (Evolinguistics: JP17H06379), and Generalitat de Catalunya (Government of Catalonia) – 2017-SGR-341.

## Data availability

The data not present in the manuscript or in the supplementary material will be available in Figshare open access repository, as well as the code used to perform the analysis reported in this study.

## Competing interests

Authors declare NO competing financial or non-financial interest.

---

**Figure S1:**
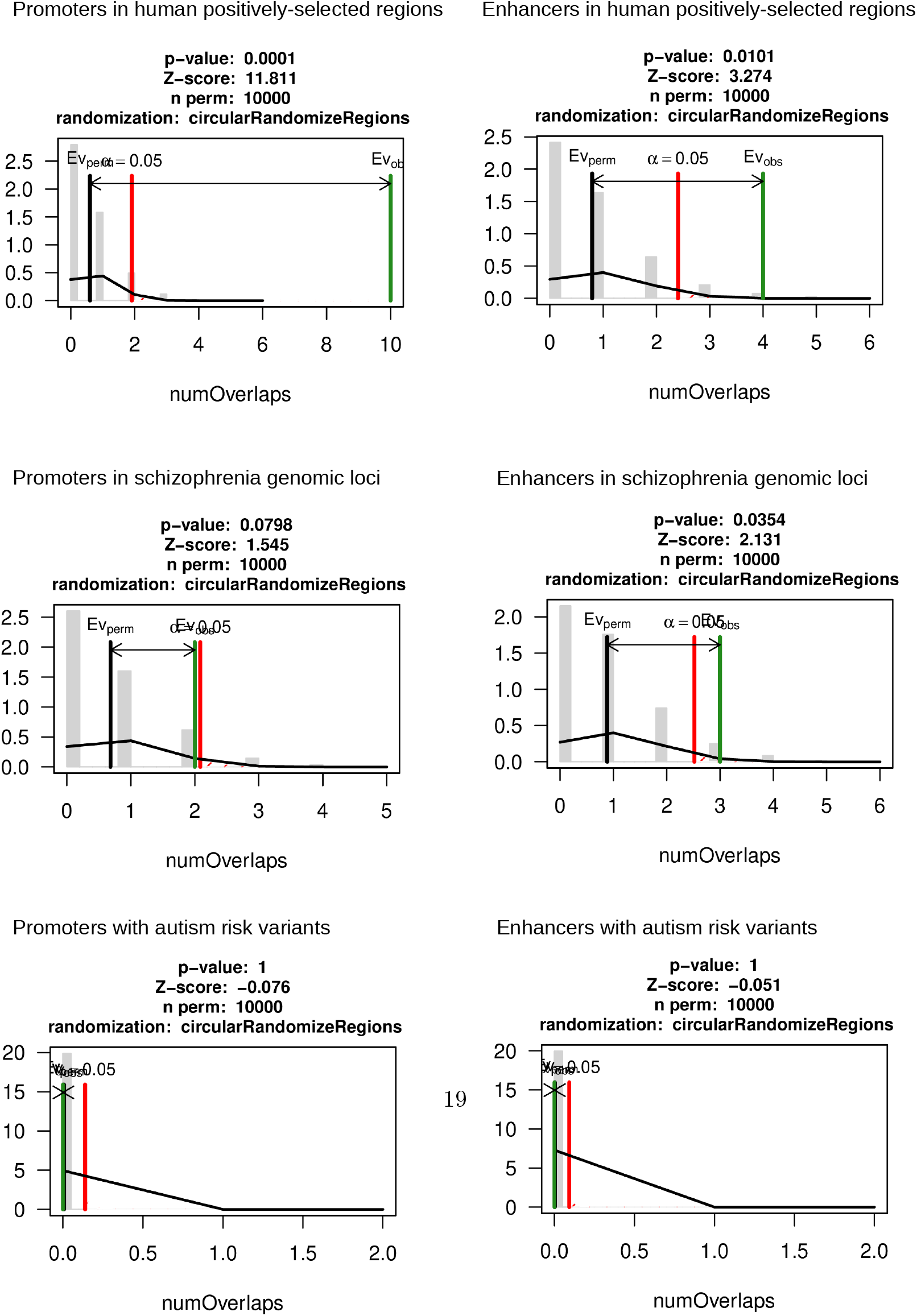
Enrichment analyses. Permutation tests were applied to determine enrichment of enhancers/promoters within human positively-selected regions (n=314, from [8]; overlap = 10 prom; 4 enh), schizophrenia genomic loci (n=108, from [20]; overlap = 2 prom; 3 enh), and autism risk variants (n=58, [21], retrieved from [22]; no overlap).

**Figure S2:**
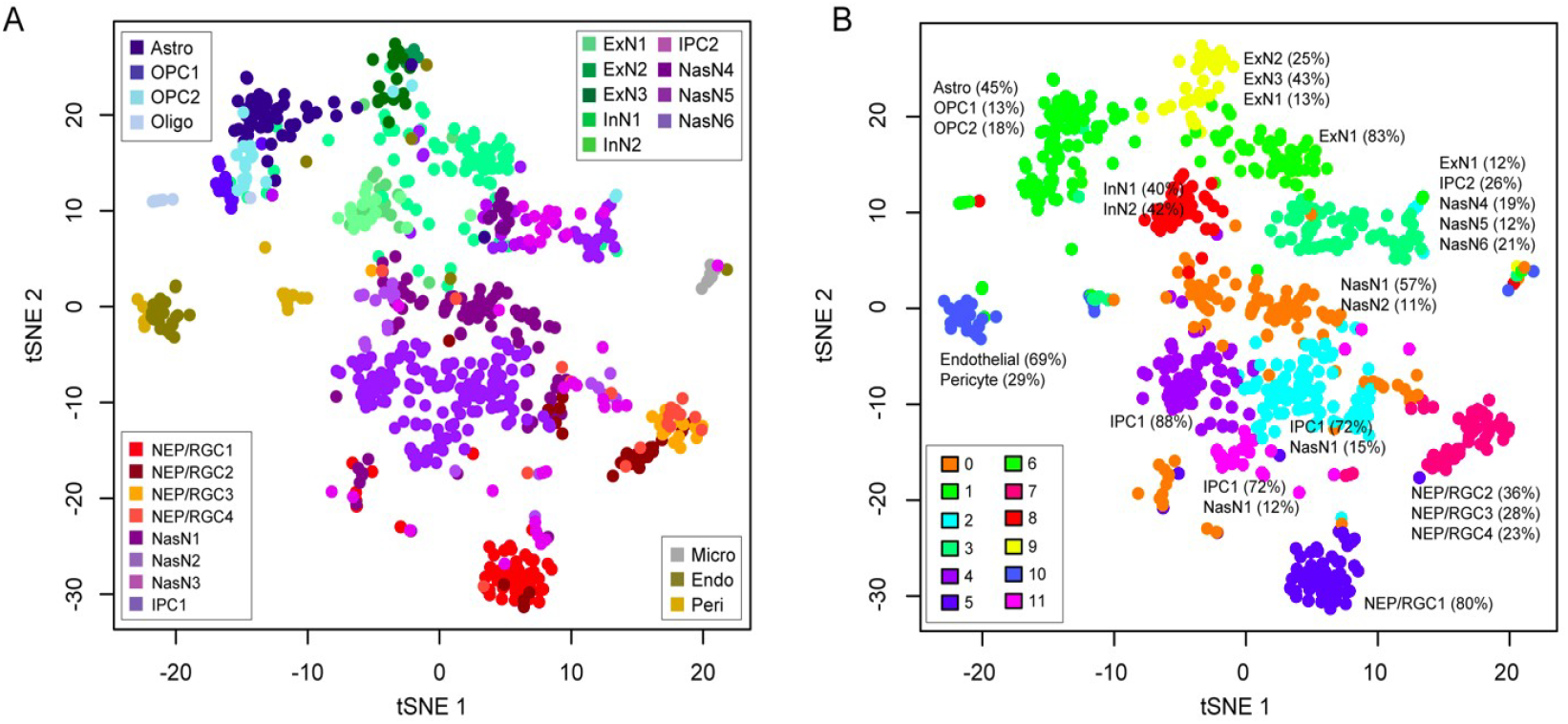
Clustering results obtained in the original study. [49] generated and analysed the single-cell transcriptomic data used in this study (5 to 20 post-conception weeks human prenatal brain). They reported two clustering results (A and B) after using different methodologies. Note in B the presence of the three sub-populations for intermediate progenitor cells and the two clusters of radial glial cells, as in our analysis. Images are reproduced with copyright permission from *Science*.

**Figure S3:**
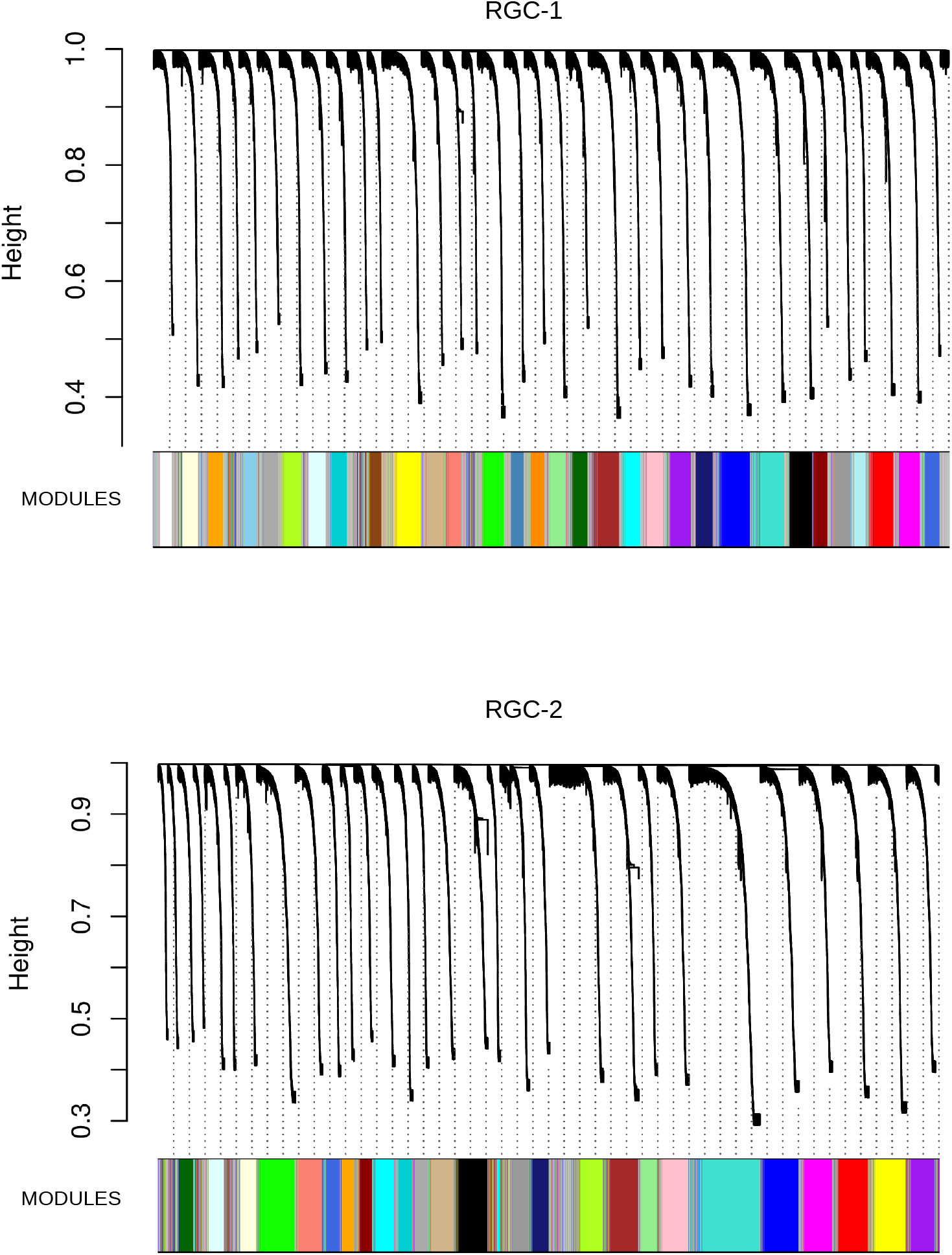
Co-expression network analysis - Radial glial cells gene modules. We used the two identified radial glial cell clusters for co-expression analyses. The dendrogram shows genes grouped in different modules (tree branches) that were assigned a color code (bottom). A total of 32 modules were detected for RGC-1 sub-population and 26 modules for RGC-2 sub-population.

**Figure S4:**
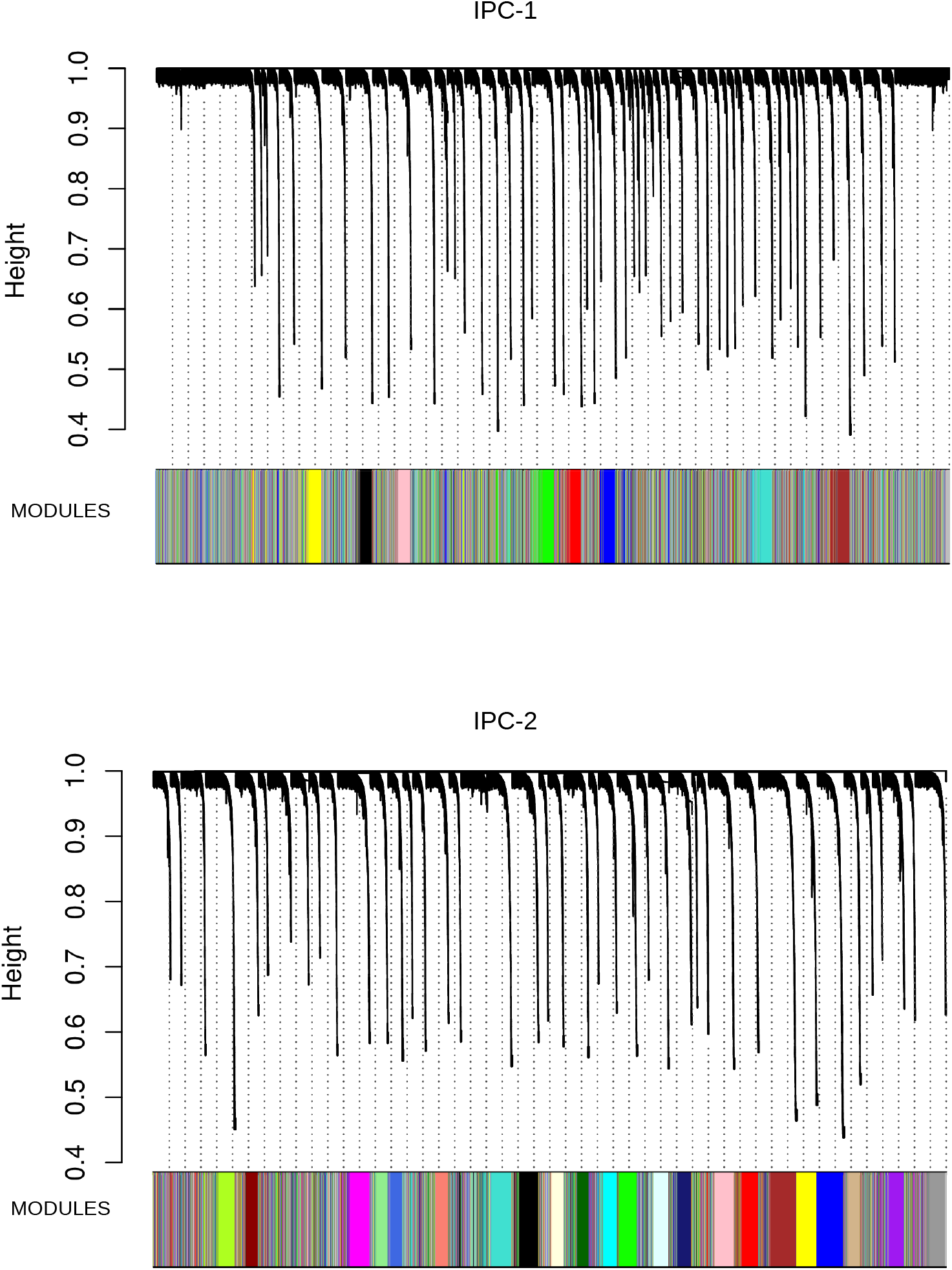
Co-expression network analysis - Intermediate progenitor cells gene modules. Two intermediate progenitor cell clusters (IPC-1 and IPC-2) were used for co-expression analyses. A total of 9 modules were detected for IPC-1 sub-population and 23 modules for IPC-2 sub-population.

